# *De novo* structural mutation rates and gamete-of-origin biases revealed through genome sequencing of 2,396 families

**DOI:** 10.1101/2020.10.06.329011

**Authors:** Jonathan R. Belyeu, Harrison Brand, Harold Wang, Xuefang Zhao, Brent S. Pedersen, Julie Feusier, Meenal Gupta, Thomas J. Nicholas, Lisa Baird, Bernie Devlin, Stephan J. Sanders, Lynn B. Jorde, Michael E. Talkowski, Aaron R. Quinlan

## Abstract

Each human genome includes *de novo* mutations that arose during gametogenesis. While these germline mutations represent a fundamental source of new genetic diversity, they can also create deleterious alleles that impact fitness. The germline mutation rate for single nucleotide variants and factors that significantly influence this rate, such as parental age, are now well established. However, far less is known about the frequency, distribution, and features that impact *de novo* structural mutations. We report a large, family-based study of germline mutations, excluding aneuploidy, that affect genome structure among 572 genomes from 33 families in a multigenerational CEPH-Utah cohort and 2,363 cases of non-familial autism spectrum disorder (ASD), 1,938 unaffected siblings, and both parents (9,599 genomes in total). We find that *de novo* structural mutations detected by alignment-based, short-read WGS occurred at an overall rate of at least 0.160 events per genome in unaffected individuals and was significantly higher (0.206 per genome) in ASD cases. In both probands and unaffected samples, nearly 73% of *de novo* structural mutations arose in paternal gametes, and predict most *de novo* structural mutations to be caused by mutational mechanisms that do not require sequence homology. After multiple testing correction we did not observe a statistically significant correlation between parental age and the rate of *de novo* structural variation in offspring. These results highlight that a spectrum of mutational mechanisms contribute to germline structural mutations, and that these mechanisms likely have markedly different rates and selective pressures than those leading to point mutations.

## Introduction

Several mechanisms, including replication infidelity (Arana and Kunkel 2010; Halliday and Glickman 1991; Keohavong and Thilly 1989), genomic damage (Friedberg 2003; Cooke et al. 2003; Marnett 2000), non-allelic recombination (Stankiewicz and Lupski 2002), and double-strand break repair (Kanaar et al. 1998) are known to create *de novo* mutations (DNMs) in the human germline. These mutations contribute to genomic diversity and often are primary targets in the analysis of rare, dominant genetic disorders. There is therefore a longstanding interest in understanding the frequency at which *de novo* mutations occur and the patterns that affect these rates. Numerous studies have measured the rate of germline *de novo* single-nucleotide and small insertions/deletions mutations (SNVs and INDELs) at approximately 70 events per individual (Sasani et al. 2019; An et al. 2018; Werling et al. 2018; Jónsson et al. 2017; Kong et al. 2012), and it has been established that the majority of these small point mutations arise on the paternal gamete. The frequency of single-nucleotide and insertion-deletion DNMs increase with parental age, especially paternal age (Sasani et al. 2019; Jónsson et al. 2017; Goldmann et al. 2016; Shendure and Akey 2015; Scally and Durbin 2012; Roach et al. 2010; Turner et al. 2017).

In contrast, precise estimates of germline mutations affecting the structure of the human genome (structural variants, or SVs) have been far more difficult to discern. *De novo* SVs (dnSVs) largely arise from mutational mechanisms that are distinct from those responsible for point mutations. The larger size of SVs increases the likelihood that any given SV will impact protein-coding genes or other critical genomic regions. Understanding the selective constraints on dnSV-specific mechanisms is essential because a broad spectrum of balanced, unbalanced, and complex structural mutations are known to underlie many developmental disorders (Ma et al. 2017; Redin et al. 2017; Chiang et al. 2012; Talkowski et al. 2012; Xu et al. 2008; Stefansson et al. 2008). However, dnSVs are predicted to occur several hundred-fold less frequently than point mutations (Werling et al. 2018), requiring a much larger sample size to achieve accurate estimates of dnSV rates.

The inherent challenge of accurately identifying SVs further complicates the measurement of dnSV rates. The short-reads that comprise most large whole-genome sequencing (WGS) datasets yield high false positive and false negative rates (Cameron et al. 2019; Mahmoud et al. 2019; Kosugi et al. 2019; Chaisson et al. 2019), as paired-end short-read alignments cannot always reveal the complete structure of an SV. Most SV detection algorithms (Chen et al. 2016; Kronenberg et al. 2015; Layer et al. 2014; Rausch et al. 2012) screen for clusters of split alignments and paired-end reads with discordant strand orientation or insert sizes, while SVs that alter copy number are also detectable through the changes in sequence depth for the variant region (Kloosterman et al. 2015; Klambauer et al. 2012; Abyzov et al. 2011).

Repetitive genomic regions obfuscate SV calling by creating inconsistent or inaccurate read alignments, which can cause false negatives by attenuating the alignment signals supporting true SVs. These regions can also produce a high rate of false-positive SV signals (Cameron et al. 2019). Long-read sequencing technologies and *de novo* assembly promise to address some of these challenges and greatly improve the accuracy and sensitivity of SV detection (Chaisson et al. 2019); unfortunately, they remain prohibitively expensive for most large-scale analyses. Detection of SVs in repetitive regions is critical for understanding dnSV rates, as multiple homology-mediated mechanisms have been shown to drive SV formation, including those arising from non-allelic homologous recombination (NAHR), (Liu et al. 2011; Sharp et al. 2005; Bailey et al. 2001) and those derived from mechanisms dependent on minimal sequence homology such as Fork Stalling and Template Slippage (FoSTeS, (Lee et al. 2007)) and microhomology-mediated break-induced replication (MMBIR, (Hastings et al. 2009)). Identifying these variants from short-read WGS, as well as those resulting from repair mechanisms that do not require sequence homology, such as non-homologous end joining (Lees-Miller and Meek 2003), requires exhaustive SV calling through the application of multiple algorithms (Collins et al. 2020; Werling et al. 2018) and extensive curation of SV predictions to remove false positives (Pedersen and Quinlan 2019; Belyeu et al. 2018). Despite these efforts, some SVs remain undetectable via short-read WGS (Chaisson et al. 2019). These inaccessible variants display sequence contexts distinct from those that can be captured by both short-read and long-read technologies, and estimates of dnSV rates utilizing such data are therefore lower bounds on the true SV mutation rate (Zhao et al. 2020).

Due to these complications, there is greater variance in estimates of dnSV rates than for single-nucleotide DNMs. Early studies from microarray technology estimated 1 very large (greater than 300kb) *de novo* copy number variant (CNV) occurs once in every 98 births (Sebat et al. 2007). More recent studies with higher resolution short-read WGS in trio and quartet families observed 1 dnSV per 5-6 births (Brandler et al. 2018; Werling et al. 2018), and estimates from larger population-based sequencing analyses were higher still at 1 dnSV per approximately 3.5 births (Collins et al. 2019). Differences in these estimates reflect variability in sample sizes, sequencing technologies, SV calling methodology, and approaches to estimating dnSV mutation rates (e.g., direct observation vs. estimations using principles of population genetics), which highlights the challenges inherent to establishing precise estimates of a human dnSV rate from short-read WGS technologies.

Estimating the rate of mutations arising from mobile element insertions (MEIs) that are still active in the human genome has also been challenging. MEI mutation rates are important to understand given their ability to impact human phenotypes by creating CNVs through non-allelic homologous recombination (Xing et al. 2009) or by interrupting genes through retrotransposition (Kazazian and Moran 2017; Hancks and Kazazian 2016). A recent study in a cohort of 33 large families identified 26 *de novo* MEI events (Feusier et al. 2019) and estimated L1 and SVA retrotransposition rates at about 1/63 births, while the rate of *Alu*Y retrotranspositions was measured at about 1/40 births. Additional investigation of *de novo* MEIs with larger cohort sizes and more MEIs will help to refine these estimates.

*De novo* structural variants are known to play a role in the genetic etiology of sporadic ASD, and multiple previous studies have shown that simplex ASD cases are more likely to harbor very large dnSVs than their unaffected siblings or the general population (Brandler et al. 2016; Sanders et al. 2011; Yoon et al. 2009; Sebat et al. 2007). Furthermore, while parent-of-origin and parental age effects have been observed for single nucleotide *de novo* mutations, their impact on dnSV rates remains unclear. Prior studies have indicated a large paternal contribution to dnSVs (Kloosterman et al. 2015), while a maternal bias was reported for a subset of recurrent dnSVs in other studies (Duyzend et al. 2016). Efforts to identify effects of parental age on dnSV rates have generally failed to show age-based enrichment but remain inconclusive due to the small numbers of dnSVs found (Girard et al. 2016; Kloosterman et al. 2015).

In this study we analyze the rate of *de novo* mutation for six major classes of SVs: deletion (DEL), duplication (DUP), insertion (INS, including MEIs), inversion (INV), translocation (CTX), and complex variants that combine more than one of the previous (CPX). By studying the genomes of a large cohort of nuclear families, we provide an accurate, lower-bound measure of the rate of *de novo* SV mutation detectable with short-read WGS data. We also explore the effects of gamete-of-origin and parental age on dnSVs and investigate potential rate differences between ASD cases and controls.

## Results

### Identification of dnSVs

Our analyses focused on two family-based cohorts with short-read WGS. The first cohort consisted of 572 samples in 33 large three-generation families from the CEPH-Utah cohort (Dausset et al. 1990); in total, these families are comprised of 434 distinct mother, father, child trios. The second cohort consisted of 2,384 families from the Simons Foundation Autism Research Initiative (SFARI) Simons Simplex Collection (SSC, (Fischbach and Lord 2010)). The SSC cohort includes 443 ASD “trios” (consisting of one affected child and two unaffected parents), and 1,941 ASD “quartets” (consisting of one affected child, one unaffected child, and two unaffected parents). We excluded samples which failed quality control analysis (see Methods), resulting in a final cohort of 2,363 ASD probands and 1,938 siblings, all with both parents available. Families selected had no known history of ASD, increasing the likelihood that SVs contributing to ASD arose *de novo* in a gamete transmitted to the affected child.

We applied a comprehensive suite of SV identification algorithms, consisting of Lumpy (Layer et al. 2014), Manta (Chen et al. 2016), Delly (Rausch et al. 2012), Whamg (Kronenberg et al. 2015), CNVnator (Abyzov et al. 2011), cn.MOPS (Klambauer et al. 2012), and MELT (Gardner et al. 2017), the latter six as part of the GATK-SV framework (Collins et al. 2020; Werling et al. 2018). We then filtered putative dnSVs using depth-of-coverage (Pedersen and Quinlan 2019) and visual inspection (Belyeu et al. 2018; Robinson et al. 2011), resulting in a set of 804 high-confidence dnSVs, excluding trisomies and sex chromosome anomalies (**Figure 1**).

**Figure 1.**
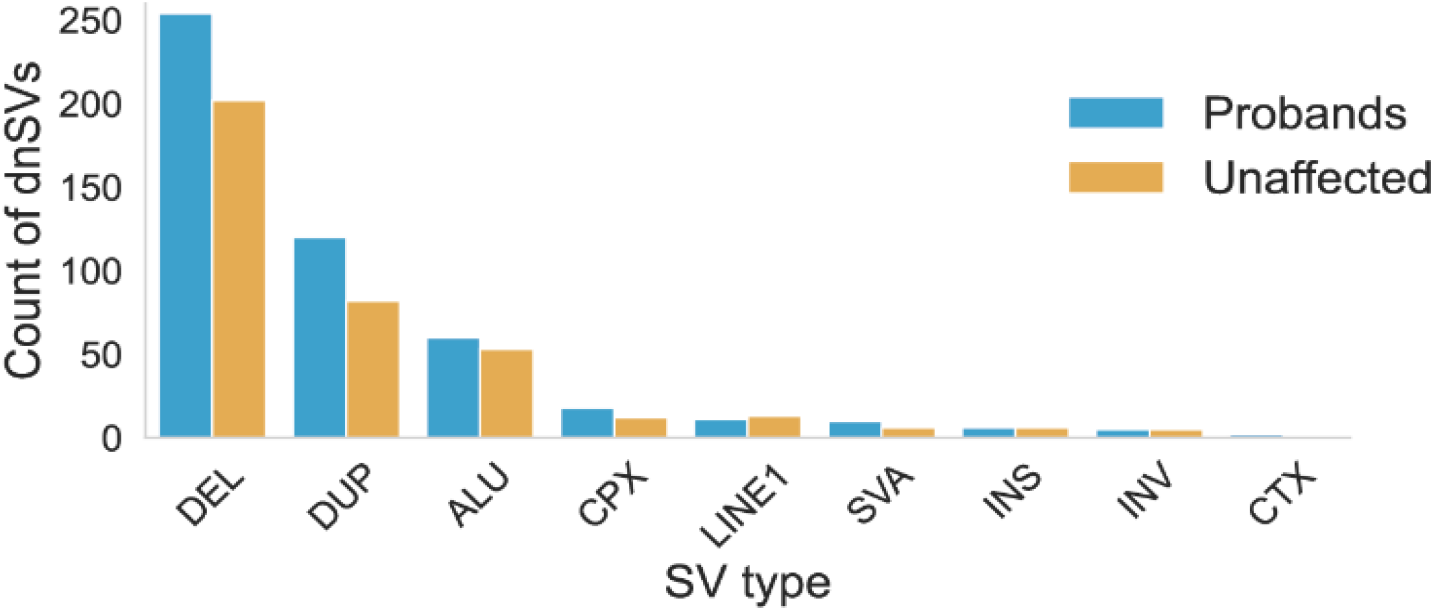
*De novo* structural variants identified in SFARI SSC. The count of dnSVs of each SV type found in 2,363 ASD probands and 2,372 unaffected samples.

Although a much smaller cohort, the unique, three-generation composition of the CEPH-Utah families enables direct measurement of the rate of false positive dnSV calls. Since these families have a median of eight (min = 4, max = 16) offspring in the third generation, any dnSV detected in a sample from the second generation of a CEPH-Utah family should have a 50% probability of transmission to each third-generation child. For example, the chance of at least one child inheriting a *true* dnSV from the second-generation parent is typically over 99% (e.g., 1-0.5^8 for a family with 8 children in the third generation). Thus, any predicted dnSV observed in a second-generation individual that is absent from all third-generation offspring is considered a false positive. We identified eight second-generation dnSVs in the CEPH families, all of which were transmitted to at least 1 third-generation offspring. Furthermore, very similar dnSV discovery methods were used in our prior study of a smaller SSC cohort, resulting in a 97% molecular validation rate of all high-quality dnSV predictions (Werling et al. 2018). Together these suggest a low false discovery rate in CEPH as well as SFARI.

Another potential problem in dnSV detection is a false negative SV call in a parent sample, which leads to an inherited SV in the child being falsely labeled as a *de novo* mutation. To test for this, we also called dnSVs in the third generation of the CEPH cohort, resulting in 54 putative dnSVs. We then visually scrutinized the evidence in the parents and grandparents of each third-generation CEPH sample with a dnSV call. We examined IGV and Samplot (Belyeu et al. 2020) images that included the offspring sample, both parents, and both sets of grandparents and carefully examined each for any missed split read, discordant pair, or coverage depth signals indicating that the putative dnSV was actually a missed transmission event. This provided an extra opportunity to detect elusive inherited variants. In 53 of the third-generation CEPH dnSVs, no evidence was detected in parents or grandparents to support the variant, while one variant presented a complex breakpoint pattern in one parent and one grandparent which might have resulted in the third-generation dnSV call (**Supplemental Figure 1**). We therefore estimate that ∼2% of variants identified as dnSVs could be cases of missed transmission.

### Analysis of *de novo* SV rates and gamete of origin

We combined the SSC and CEPH dnSV sets, yielding a final set of 865 dnSVs in 4,735 offspring genomes (9,599 genomes with parents included), including 2,363 ASD probands and 2,372 unaffected offspring.

Prior studies involving fewer families have shown that ASD cases are often enriched for dnSVs compared to unaffected siblings (Brandler et al. 2016; Sanders et al. 2011; Yoon et al. 2009; Sebat et al. 2007). One study (Werling et al. 2018) analyzed WGS in 519 SFARI families that were pre-screened for *de novo* loss-of-function variants or large CNVs and found no enrichment of dnSVs in probands. We leveraged this now much larger set of SSC and CEPH families to test the dnSV rate for probands and unaffected controls (**Figure 2A)**. We found a statistically significant increase in dnSVs among ASD probands (486 dnSVs / 2,363 cases) compared to unaffected samples (379 dnSVs / 2,372 unaffected samples; p=0.0008, Fisher’s Exact Test). The rate of dnSVs in this cohort was therefore one mutation for every 0.2056 births in probands and one mutation for every 0.1598 births in controls. This enrichment was significant for duplications (p=0.0212, **Supplemental Figure 2A**), yet not significant for deletions (p=0.0556, **Supplemental Figure 2B**) or *Alu*-family MEIs (p=0.7036, **Supplemental Figure 2C**).We developed a method to use informative SNVs within or near dnSVs to determine the parental gamete-of-origin (the mutation’s haplotype phase) for *de novo* deletions, duplications, inversions, and complex variants. 257 dnSVs successfully phased, including 36.8% of all dnSVs of those types, see Methods for additional details. This analysis revealed an enrichment for paternally derived SVs in both probands and unaffected samples (**Figure 2B**). Among unaffected samples, 62 (69.7%) dnSVs arose on the paternal gamete and 27 (30.3%) from the maternal gamete; these rates were significantly different (Fisher’s Exact test; p=0.0026). Similarly, among probands, 125 (74.4%) dnSVs had a paternal origin, and 43 (25.6%) were maternally derived; the difference in these rates was also statistically significant (p<0.0001, Fisher’s Exact Test).

**Figure 2.**
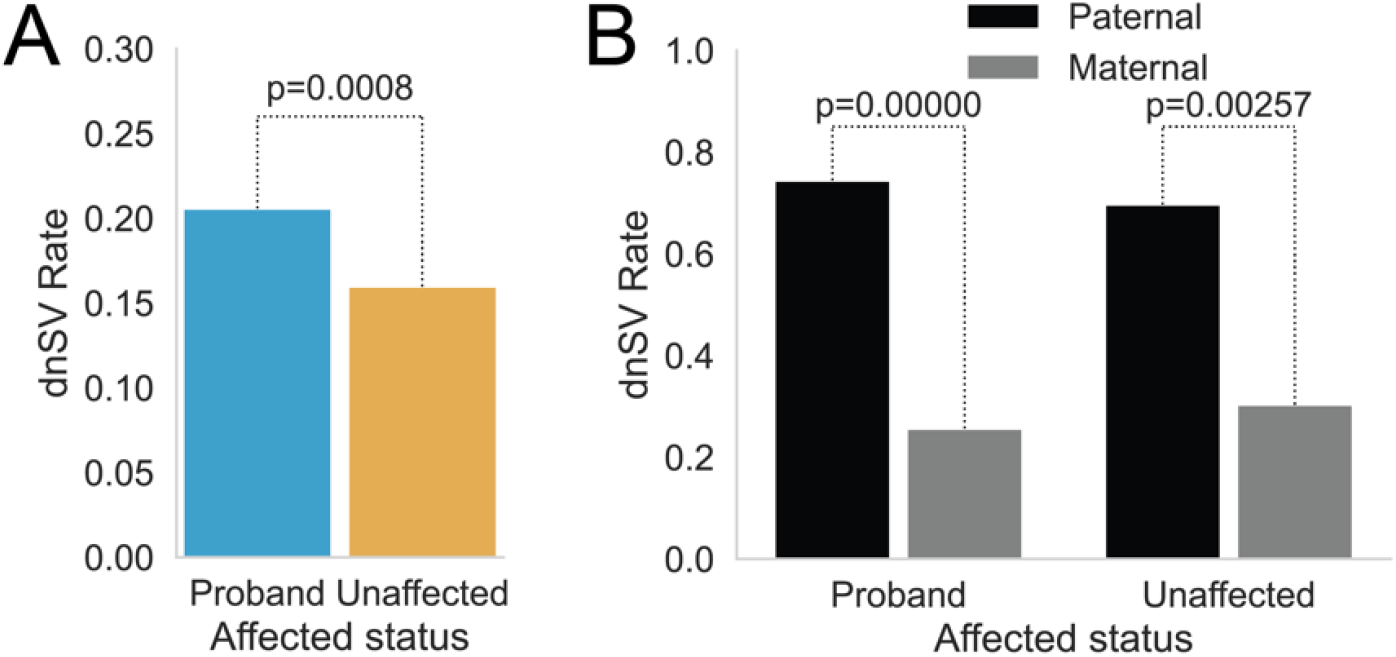
Comparisons of dnSV rates. **A**. Comparison by Fisher’s Exact test of dnSV rates in probands vs. siblings. **B**. Comparison by Fisher’s Exact test of dnSV rates from variants phased to maternal of paternal gamete in probands and siblings.

### Age effects on dnSV rates

An important unanswered question is whether dnSV rates increase with parental age. Parental age effects on the rate of *de novo* single-nucleotide mutations have been previously identified and are an important factor in profiling the likely causes of genetic disease in individuals with older parents. A goal of our study was therefore to determine if there is a similar increase in the rate of dnSVs with parental age. We used the father’s age at birth of the child as a proxy for parental age and grouped samples by dnSV status (0 vs. 1 or more dnSVs), then performed a one-sided Wilcoxon rank-sum test for an increase in paternal age among samples with a dnSV versus those without (**Figure 3**). In probands we found no significant difference in the distributions of father’s ages between the two groups (p=0.554), while in unaffected samples we found a significant increase in father’s ages (p=0.033) which did not remain significant after Bonferroni multiple test correction for two tests (adjusted p=0.066). We estimate that we have 80% power to detect a mean paternal age difference of 0.851 and 0.940 years between samples having a dnSV versus those without a dnSV in probands and unaffected samples, respectively (**Supplemental Figure 3**). Thus, while undetectable within our cohort, a parental age bias may still exist, albeit with a much weaker effect than observed for some other types of mutation such as *de novo* SNVs; detecting a statistically significant age effect will likely require an even larger cohort. A potential confounding variable in this comparison is the known effect of paternal age on risk for ASD (Reichenberg et al. 2006), which could act to decrease our power to detect a parental age effect in ASD probands.

**Figure 3.**
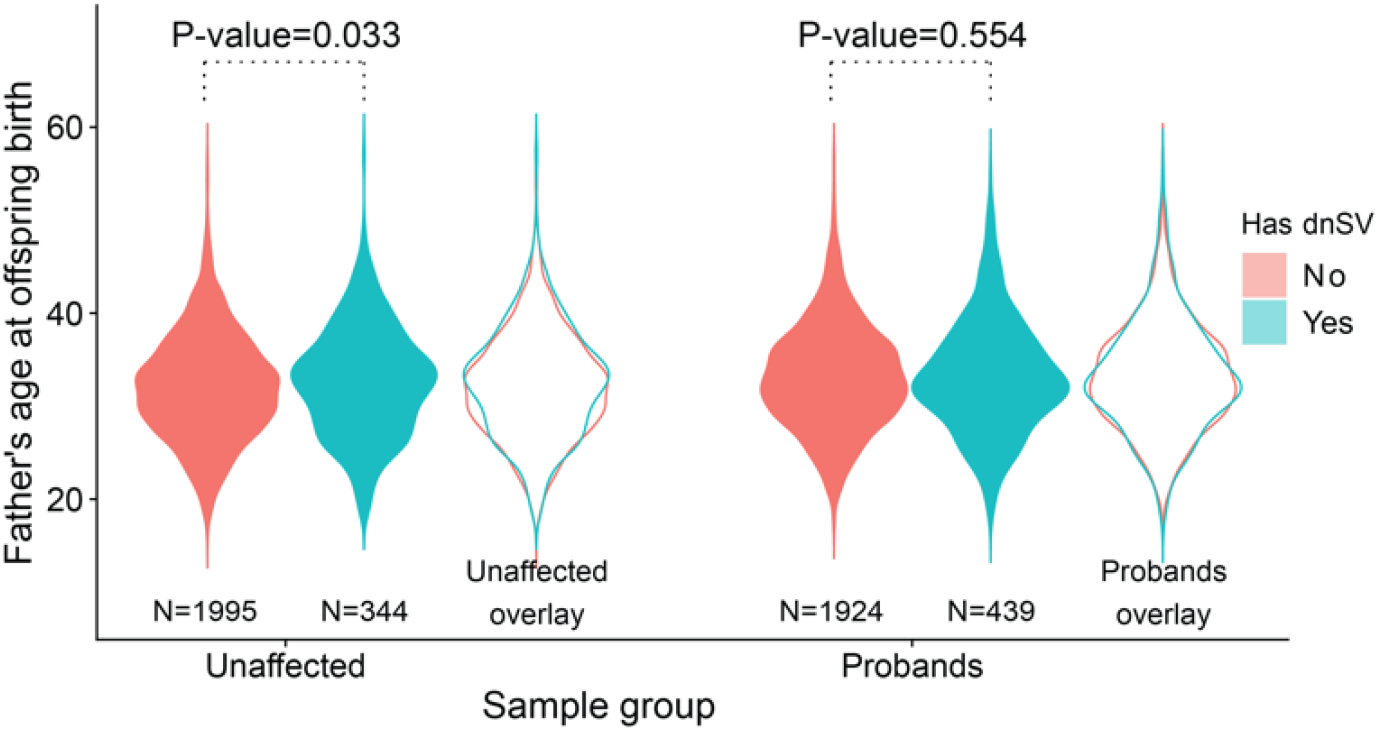
Correlations of paternal age and *de novo* structural variant rate. Comparison by one-sided Wilcoxon rank-sum test for increased father’s age in samples with at least one dnSV versus those without. This test is performed for unaffected samples (left) and probands (right). Overlay highlights differences in father’s age distribution between samples with and samples without a dnSV. After Bonferroni multiple test for two tests, the adjusted marginal significance for unaffected samples is p=0.066, and for probands the adjusted marginal significance is p=1.

We tested for effects of either paternal or maternal age using the subset of dnSVs which had been phased to a parental gamete-of-origin (165 probands with dnSVs, 85 unaffected samples with dnSVs) and found no difference in parental ages between samples with or without a dnSV (one-sided Wilcoxon rank-sum test, **Supplemental Figure 4**). We hypothesize that this lack of correlation speaks to the inherently different mechanisms underlying point mutations and structural changes, as the increase in point mutation rate with parental age is substantial and reproducible. As a control, we also performed a Poisson regression to test the effect of paternal age on the rate of *de novo* single-nucleotide mutations in this cohort and, as expected given prior findings (Sasani et al. 2019; Jónsson et al. 2017), we found a significant correlation (p<2e-16) between the number of *de novo* SNVs and the age of the father for both unaffected samples and probands (**Supplemental Figure 5**).

Next, we tested for parental age effects on rates of the most common SV types: deletions (254 in probands, 202 in unaffected samples), duplications (120 in probands, 82 in unaffected samples), and MEIs (87 in probands, 78 in unaffected samples). We used father’s age as a proxy for parental age and found no enrichment with paternal age in deletions or duplications, although there was an increase in dnMEI risk to unaffected samples with increased paternal age (difference in means 2.18 years, p=0.004, not significant after Bonferroni correction for eight tests, **Supplemental Figure 6**). The dnMEI age effect may therefore drive the signal detected in analysis of all SV types shown in Figure 3, as neither of the other common SV types showed a significant enrichment. This may reflect the fundamentally different mechanisms underlying dnMEIs and argues for future research, especially with long-read sequencing technologies that offer greater power to characterize mobile element insertions.

### Mechanisms responsible for dnSVs

Identifying the primary mechanisms responsible for dnSVs is of fundamental interest to characterizing mutational hotspots and to understanding the mutational forces driving evolution of genome structure. However, inferring the exact mechanism underlying each SV is complicated by imprecise breakpoint mapping and low sequence complexity at SV breakpoints. We therefore grouped variants into three categories based on the degree of sequence homology observed at each dnSV breakpoint, an essential feature for several known mechanisms of structural variation and analyzed mechanistic correlations with parental origin in the 257 phased variants (**Figure 4A**).

**Figure 4.**
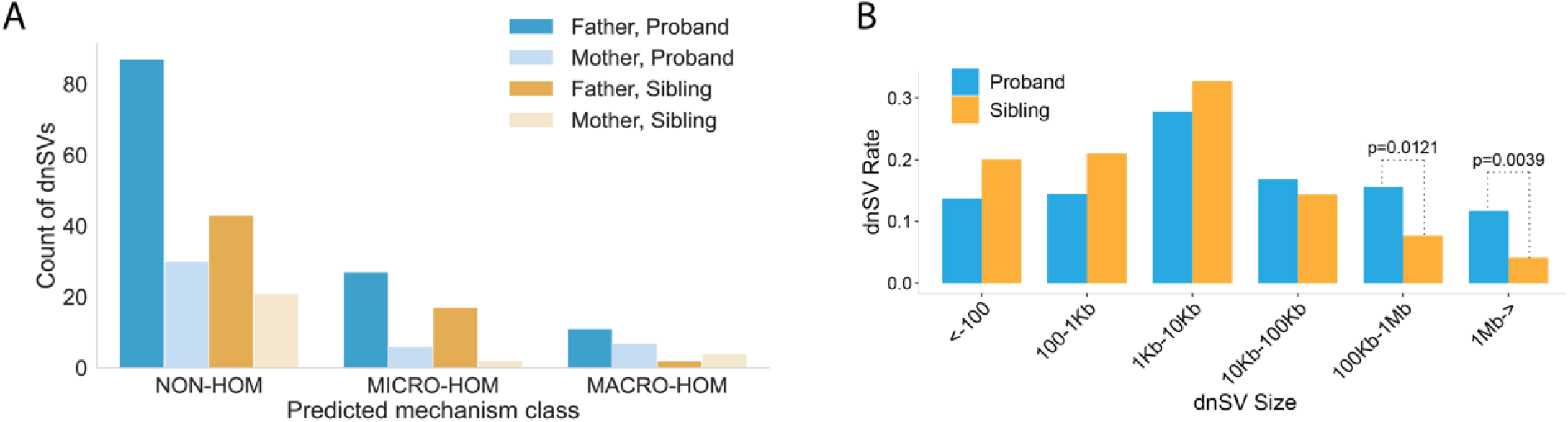
A comparison of dnSV breakpoint homology and size among probands and unaffected samples. **A**. Counts of phased variants grouped by predicted mechanism class, parent-of-origin, and affected status. Mechanism classes include those characterized by no sequence homology at breakpoints (NON-HOM), microhomology at breakpoints (MICRO-HOM), or macrohomology (matching segmental duplications) at breakpoints (MACRO-HOM). **B**. Variants binned by size and compared between probands and unaffected samples. The fraction of dnSVs assigned to each bin is statistically similar except in the largest two bins where sizes are 100Kb to 1Mb and >= 1Mb. The difficulty of determining size of insertion variants, especially mobile element insertions, lead to exclusion of those variants from this figure.

We found that CNVs flanked by segmental duplications, which are potential substrates for NAHR (Liu et al. 2011; Sharp et al. 2005; Bailey et al. 2001)), were grouped together in the macrohomology **(MACRO-HOM**) class. Small NAHR variants are difficult to detect, as the flanking homology at the breakpoints can decrease paired-end read signals and depth-of-coverage fluctuations that are used by Illumina-based SV-detection methods to identify variants, but large CNVs derived from NAHR are readily detected; we identified 45 NAHR-derived mutations with a median length of 1.5 megabases. Various mechanisms of SV are identifiable by short sequences of breakpoint-flanking microhomologies, including fork stalling and template switching (FoSTeS, (Lee et al. 2007)) and microhomology-mediated break-induced replication (MMBIR, (Hastings et al. 2009)). We therefore grouped dnSVs with a 2-10bp homologous sequence flanking breakpoints as the microhomology (**MICRO-HOM**) class. Finally, we grouped variants with no breakpoint homology, likely resulting from incorrect joining of double-strand breaks in many cases (Lees-Miller and Meek 2003), as the **NON-HOM** (non-homology) class. Only two variants that we could capture from short-read WGS had breakpoint homology outside these size categories (18 and 22bp).

We categorized all dnSVs except MEIs, for a total of 707 events, by the extent of breakpoint homology and inferred the mechanistic class most likely responsible (see Methods for details). The majority of dnSVs (530, ∼75%) lacked sequence homology, while fewer exhibited either macro-homology (45, ∼6%) or micro-homology (132, ∼19%). More dnSVs derived from the paternal gamete in each mechanism class. There were similar rates of micro-homology and non-homology dnSVs in probands and unaffected samples, but a higher rate of macro-homology dnSVs in probands (N=36 vs N=9). This may relate to the large size of those variants (1.5 megabases), as compared to the 5.4 kilobase median length of all deletions and duplications we discovered. This substantially larger variant size also increases the risk of impacting genes or regulatory elements potentially involved in development of ASD. As 257 or about 36% of these dnSVs were successfully phased, the analysis of parental effect on mechanism includes 24 macrohomology dnSVs, 52 microhomology dnSVs, and 181 non-homology dnSVs (**Figure 4A**).

Many insertion variants were identified as MEIs. The active classes of MEIs consist of L1, *Alu*, and SVA retrotransposable elements (**Figure 1**). We identified 110 *de novo Alu* events, 15 *de novo* SVA events, and 20 *de novo* L1 events in probands and unaffected samples, for approximate rates of 1 *Alu* per 42 births, 1 SVA per 309 births, and 1 L1 per 231 births. These rates of *Alu*-family DNMs are quite similar to a recently reported measurement of 1 *Alu* per 40 births in the CEPH cohort (Feusier et al. 2019), while LINE1 and SVA rate incongruity (each reported in that study as 1/63 births) could result from the approximate ten-fold difference in cohort size, our exclusion of somatic mosaic events (which were included in the Feusier et al. 2019 rates), or from use of three independent MEI detection software tools in that study to achieve extremely high variant recovery. MEI rates were similar between probands and unaffected samples.

Finally, we compared the sizes of *de novo* deletions, duplications, and inversions found in ASD probands and unaffected samples. After Benjamini-Hochberg multiple test correction (with alpha=0.05), we found no significant differences except for the two largest size bins, containing dnSVs impacting at least 100Kb. Such large dnSVs were observed in 112 probands compared to 38 unaffected samples (**Figure 4B)**. This enrichment reflects the known role of large genomic alterations in ASD (Brandler et al. 2016; Sanders et al. 2011; Yoon et al. 2009; Sebat et al. 2007).

## Discussion

To our knowledge, this is the largest direct measurement of dnSV from family-based WGS to date. Our results demonstrate that parent of origin is the predominant factor influencing the frequency of *de novo* SV events. We also reproduce the well-established result that the *de novo* rates of SVs and single-nucleotide mutations are higher in ASD probands than in unaffected samples. While there is consensus that the burden of single-nucleotide and insertion-deletion germline mutations increases with parental age, we find that any correlation between parental age and the burden of *de novo* structural variation in this study must be modest if present at all. This result suggests that fundamentally different endogenous and exogenous mechanisms create *de novo* point mutations versus structural mutations, and our results suggest an increased negative selective pressure on large chromosomal rearrangements.

We discovered large NAHR events (N=45; median size = 1.5 Mb) identified by the presence of large, high-identity segmental duplications flanking the breakpoints of the dnSV. The size of these mutations may explain much of the imbalance between rates in ASD proband and unaffected samples of potential NAHR-derived variants, since larger variants are more likely to be deleterious and are known to be under stronger negative selection (Collins et al. 2019; Sudmant et al. 2015). Identification of repeat-flanked variants is also quite difficult in some cases, especially in small regions, so smaller macrohomology-mediated SVs likely were not detected. The extensive high-identity breakpoint homologies that are required for NAHR increase the difficulty of mapping and alignment, greatly decreasing the signal of discordant pairs and split reads used by most SV detection algorithms. Smaller variants caused by NAHR are also likely to be missed by depth-based SV calling tools as the deviations in depth of coverage are often too small for confident assessment of copy number status. Our discovery of a mechanism bias in these ASD cases is therefore most probably driven by size, rather than mechanism.

We note that we identified 63 copy number variants in the SSC cohort and two in the CEPH cohort that appear to be somatic in origin and mosaic in blood cells. These mutations had strong discordant and split-read alignment evidence but little to no impact on depth of coverage, likely reflecting their low cellular prevalence. A similar number of blood-mosaic mutations were observed in probands (N=35) and unaffected samples (N=30). Since these mutations did not arise in the germline, they were excluded from our mutation rate analyses. However, depending on when these mutations arose in development, some may be transmittable to the next generation if they arose prior to the establishment of the individual’s germ cell lineage.

Overall, our analysis of DNMs in over 2,300 families has established confident lower bound estimates for the rate of SV mutation in the human germline and has quantified the effects of parental age on dnSV risk, as well the mechanisms underlying dnSVs. While studying the genomes of more than 4,300 offspring and their parents provides substantial power to estimate *de novo* structural mutation rates, we emphasize that our estimates are lower bounds. While SV detection with short, paired-end WGS has improved dramatically over the last decade, sensitivity remains a challenge, especially for smaller SVs, insertions, and repeat-mediated SVs. Recent comparisons of SVs detected with short-reads and long-read technologies, such as PacBio or Oxford Nanopore Technologies, found thousands of SVs were missed by short-read technologies (Chaisson et al. 2019), and quantified the relative impact of sequence context on detection rates. Therefore, we anticipate that future studies of dnSV based upon long-reads will increase the detection of dnSVs, especially for smaller mutations arising in tandem-repeat sequences that are known to be hypermutable (Chakraborty et al. 1997).

## Methods

### *De novo* structural variant identification

We detected SVs in the CEPH and SFARI cohorts with the GATK-SV discovery pipeline previously described (https://github.com/broadinstitute/gatk-sv (Collins et al. 2020; Werling et al. 2018)). Briefly, GATK-SV is an ensemble approach that uses multiple established SV detection tools to maximize sensitivity while re-evaluating evidence directly from BAMs using a random forest classifier to improve specificity in a series of variant classification modules. GATK-SV has flexibility with initial input SV algorithms, and in this case we ran an amalgamated series of complementary algorithms that detect SVs based on a variety of signatures including discordant paired-end reads (PE), split reads (SR), and read depth (Delly (v0.7.8), smoove (v0.2.4), Manta (v1.3.1), Wham (v1.7.0) and MELT (v2.1.4), CNVnator (v0.3.3), and a custom version of cn.MOPS). Upon completion of the pipeline a VCF is derived containing adjudicated and integrated SVs from the raw algorithms. Furthermore, complex SVs (Brand et al. 2015; Collins et al. 2017) and other SVs involving more than one breakpoint (e.g. inversions, reciprocal translocations) are fully resolved.

Akin to SNVs and indels (Francioli et al. 2017; Li et al. 2012), careful filtering is required for precise identification of *de novo* SVs. We have developed a set of post hoc filtering criteria that work from a GATK-SV VCF. We start with an unfiltered VCF and remove any unresolved breakpoints, SVs with split read support observed on only one side a breakpoint, and any CNV found to have multi-allelic copy states making the determination of parental haplotypes challenging and therefore *de novo* calling highly inaccurate. We then exclude any variant with a parental frequency of greater than 1% for the autism cohort and 10% for the CEPH study (to account for potential F1 *de novo* transmissions). Next, only variants with support in the initial raw callers are kept, removing any potential genotyping errors, and CNV copy state overlap is investigated in parents to account for the imprecise boundaries of depth-based CNV detection that could cause an inaccurate but overlapping CNV call in a parent or child. Next we apply a set of genotype quality (GQ) filters that were determined by an ROC curve analysis with a truth set derived from molecularly validated *de novo* SVs from a previous study on a smaller subset (n=2,076) of the total cohort (Werling et al. 2018) and false positives defined as novel *de novo* variants in those samples. Both child and parental GQ cutoffs were determined, the latter of which classify variants initially predicted to be *de novo* that are in fact likely inherited. Optimal GQ parameters were found for an overall GQ, as well as depth-based GQ and PE/SR GQ all derived from the genotyping step in GATK-SV and present in the final VCF. Given the lack of validated training set for CEPH and the much smaller sample size, only a simple filter of less than 30 for depth-based GQ and PE/SR GQ in parents was applied. All variants that passed these filters were manually reviewed using duphold (Pedersen and Quinlan 2019), IGV (Robinson et al. 2011), SV-Plaudit (Belyeu et al. 2018) with Samplot (Belyeu et al. 2020), and an internal R based visualization script found in GATK-SV. In order to reduce the chance of missing a variant of interest, all private variants with a passing parental GQ are included in the manual investigation. In the SFARI cohort, any sample that was found to have *de novo* events on 4 or more chromosomes and confirmed not to have chromothripsis were excluded from manual review and classified as “outlier samples” (n=14; 0.3% of children). No outliers were excluded in the CEPH cohort.

We investigated potential *de novo* mosaic events via the following steps. Using the results of our random forest filtering, we identify CNVs greater than 10 kb from both depth and paired-end/split-read based algorithms that pass the random forest filtering’s p-value threshold but not its separation threshold. After passing through a step where fragmented variants are stitched together, we use a cohort-based variant frequency estimate to identify variants that appear only once in the cohort. We initially set a variant frequency cutoff of 1% for SFARI, but upon manual review we realized this was not stringent enough, so we increased the frequency cutoff in CEPH to 0.3%. Manual curation of the passing mosaic variants is then used to identify true events as described above.

### Identifying parent-of-origin for dnSVs

Phasing identifies the parent whose gamete underwent an error leading to a spontaneous mutation. We used single-nucleotide variants which varied in state between parents and which were inherited heterozygously (informative sites) to identify localized haplotypes that derived from either the mother or the father. We developed and applied a combination of two methods:

### Extended read-based phasing

For each dnSV we selected reads that supported the variant (split reads or discordant pairs whose gap and orientation fit the variant), then used any nearby heterozygous SNV sites to associate other reads in the region to the variant haplotype or the reference haplotype, allowing us to extend our search for informative sites from the variant breakpoints. We then tested for informative site overlap, using any informative sites up to a maximum distance from the breakpoints of 5kb, as long as an overlap with haplotype-assigned reads was found. If at least one informative site was found with read overlap, variant phasing was possible.

### SNV allele balance CNV phasing

Copy number variants have a predictable effect on the allele balance of SNVs within the region of the variant. Duplications should approximately double the number of reads that come from the duplicated region, while deletions should eliminate reads from the deleted region. Thus, where the allele balance for informative sites in the region of the variant shifts, it becomes possible to determine from which parent the *de novo* event was inherited. We identified all informative sites within deletions and tested for hemizygosity, where an informative site allele that should have been inherited from one parent instead disappeared and an allele not shared by both parents was inherited. We similarly identified informative sites within duplications and identified cases where allele balance was at least 2:1, rather than the null expectation of 1:1. In some cases of large CNVs, several informative sites were identified that gave contradictory phasing results. If at least 95% of sites supported one parent as the origin, we assigned the variant to that parent; otherwise, we excluded the variant from phasing.

This combination of phasing strategies allowed us to phase 257/698 deletions, duplications, inversions, and complex variants, an improvement on phasing rates for SNVs in previous work (Sasani et al. 2019). Insertion variants proved the most difficult type to phase, as the split reads and discordant pairs generated by these variants (especially mobile element insertions, which were the bulk of the insertions in our callset) often mis-align or align to high-copy genomic repeats and are lost, removing the evidence needed to relate variants to a haplotype. We therefore did not analyze parent-of-origin effects on insertion variants.

### Predicting causal mechanisms for dnSVs

It is impossible to confidently identify the causal mechanism for many variants, as no perfect evidence exists to confirm that a specific mechanism was responsible. However, the sequence context of a variant often provides clues that can be used to determine the most likely type of candidate mechanism that could have led to a variant’s formation. Using methodology similar to a previous analysis of SV breakpoints in mouse models (Quinlan et al. 2010), we analyzed the variants in our dnSV set with respect to four broad categories of mechanism that can lead to creation of a dnSV:

### Microhomology-based variants

We grouped together mechanisms that lead to spontaneous rearrangement due to microhomology, including microhomology-mediated break-induced replication (MMBIR, (Hastings et al. 2009)) and fork stalling and template-switching (Lee et al. 2007). For each variant we collected split reads that spanned the breakpoints, requiring at least two split reads for each breakpoint. This provided strong evidence that the breakpoint coordinates were correctly identified and allowed us to test for homology of 2-100bp between the regions upstream and downstream from each breakpoint. Variants with breakpoints that had microhomology were categorized as most likely deriving from microhomology-based mechanisms. We identified 132 microhomology SVs, with 74 in probands and 58 in unaffected samples.

### Macrohomology-based mechanisms

We used annotations of known segmental duplication pairs to identify variants that likely resulted from non-allelic homologous recombination (NAHR). NAHR variants can be difficult to identify using short-read sequencing data, as lengthy high-identity repeats must flank the resulting variant. These repeats often mask the signals used to detect SVs by increasing the difficulty of read mapping and alignment and may be especially detrimental to the accurate identification of discordant pairs and split reads, which many SV calling tools rely on. Extremely large CNVs are the most likely NAHR-derived variants to be accurately detected as these can be found using depth-based CNV callers, such as CNVnator and cn.MOPS. Breakpoint resolution was inexact with these variants as they generally lacked any confident split-read support (due to flanking repeats). Thus, we grouped together CNVs whose breakpoints were flanked by segmental duplications of at least 95% sequence identity. We required that the end of the first of the pair of segmental duplication be within a distance of 20% of the CNV length from the start of the CNV, and the start of the second of the segmental duplication pair be within a distance of 20% of the CNV length from the end of the CNV, due to the extreme error in breakpoint calling that often arises in repeat-rich regions. 45 dnSVs were assigned to the NAHR category, including 36 in probands and 9 in unaffected samples.

### Non-homology-based mechanisms

Variants which were not identified as MEIs and did not have either macro or microhomology flanking the breakpoints were classified as non-homology based. These variants may arise from a number of molecular mechanisms, such as non-homologous end joining (Davis and Chen 2013)), in which double-strand breaks are corrected and filled in an error-prone manner. We grouped 530 variants as non-homology-based, including 288 in probands and 242 in unaffected samples.

## Acknowledgements

We are grateful to the families participating in the Simons Foundation Autism Research Initiative (SFARI) Simplex Collection (SSC) and the Utah individuals who participated in the CEPH consortium. Research and contributing authors were supported by the National Institutes of Health (NIH) grants: HG006693, HG009141 and GM124355 to A.R.Q; MH115957, HD081256, HG008895, HD096326 and HD099547 to M.E.T.; K99DE026824 to H.B.; GM118335 and GM059290 to L.B.J. and the Simons Foundation for Autism Research Initiative (SFARI #573206 to M.E.T. and 388196 to A.R.Q.), with additional support from the Utah Genome Project and the George S. and Dolores Doré Eccles Foundation. Dr. Talkowski was also supported by the Desmond and Ann Heathwood MGH Research Scholars award. We would like to thank the SSC principal investigators (A.L. Beaudet, R. Bernier, J. Constantino, E.H. Cook, Jr, E. Fombonne, D. Geschwind, D.E. Grice, A. Klin, D.H. Ledbetter, C. Lord, C.L. Martin, D.M. Martin, R. Maxim, J. Miles, O. Ousley, B. Peterson, J. Piggot, C. Saulnier, M.W. State, W. Stone, J.S. Sutcliffe, C.A. Walsh, and E. Wijsman) and the coordinators and staff at the SSC clinical sites; the SFARI staff, in particular N. Volfovsky; D. B. Goldstein for contributing to the experimental design; the Rutgers University Cell and DNA repository for accessing biomaterials; the New York Genome Center for generating the WGS data. We also thank Ray White, Jean-Marc Lalouel, and Mark Leppert, who were instrumental in the ascertainment of the CEPH/Utah pedigrees.

