## Supplemental figures for "*De novo* structural mutation rates and gamete-of-origin biases revealed through genome sequencing of 2,396 families"

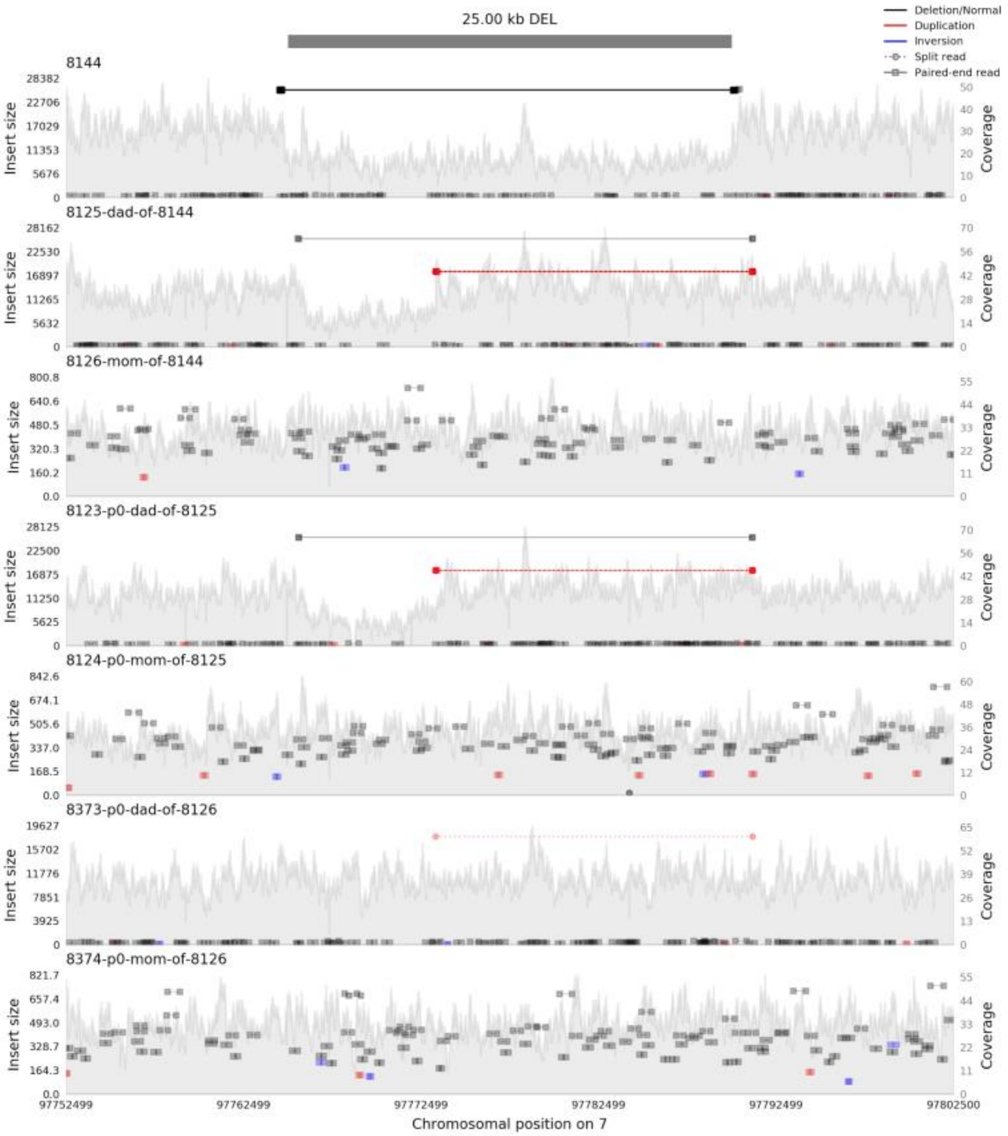


**Supplemental Figure 1. Sequence data for a possible false dnSV.** This samplot image shows paired-end reads spanning a putative *de novo* deletion in sample 8144, with a corresponding drop in coverage. The father and paternal grandfather of 8144 have a complex variant signal in a similar region with slightly different coordinates, indicating that the deletion variant could be a partial transmission of the complex variant or an unrelated *de novo* event

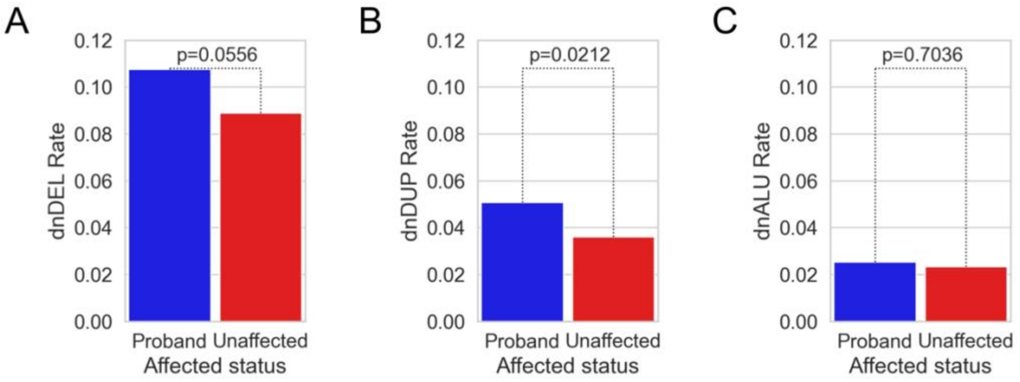


**Supplemental Figure 2. Comparison of rates of de novo structural variation by SV type. A.** No significant enrichment for de novo deletions in probands vs. siblings. **B**. Significant enrichment for de novo duplications in probands vs. siblings. **C**. No significant difference between *Alu* rates in probands vs. siblings.

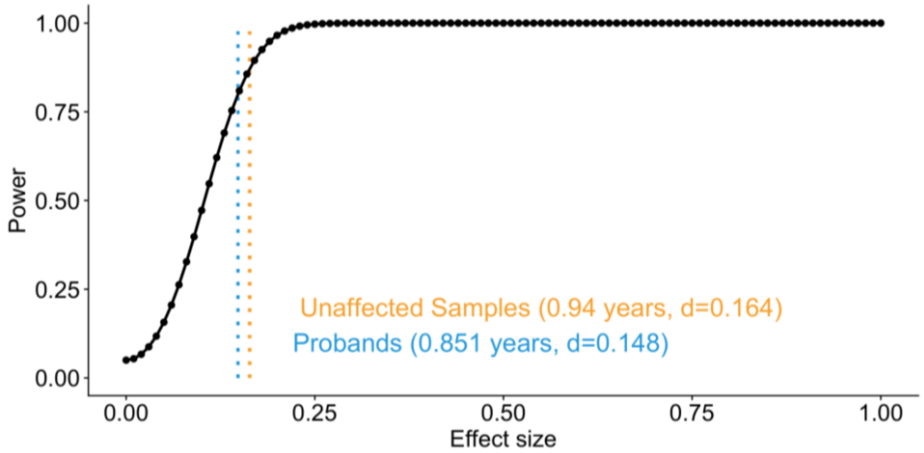


**Supplemental Figure 3. Analysis of power to detect a paternal age effect on dnSV rate.** Cohen's d statistic was used as a measure of effect size. Dotted vertical lines indicate the minimum effect size detectable at power=0.8 for each group. The effect size, d, is given in difference in number of pooled standard deviations between means and in number of years difference in father's age between means.

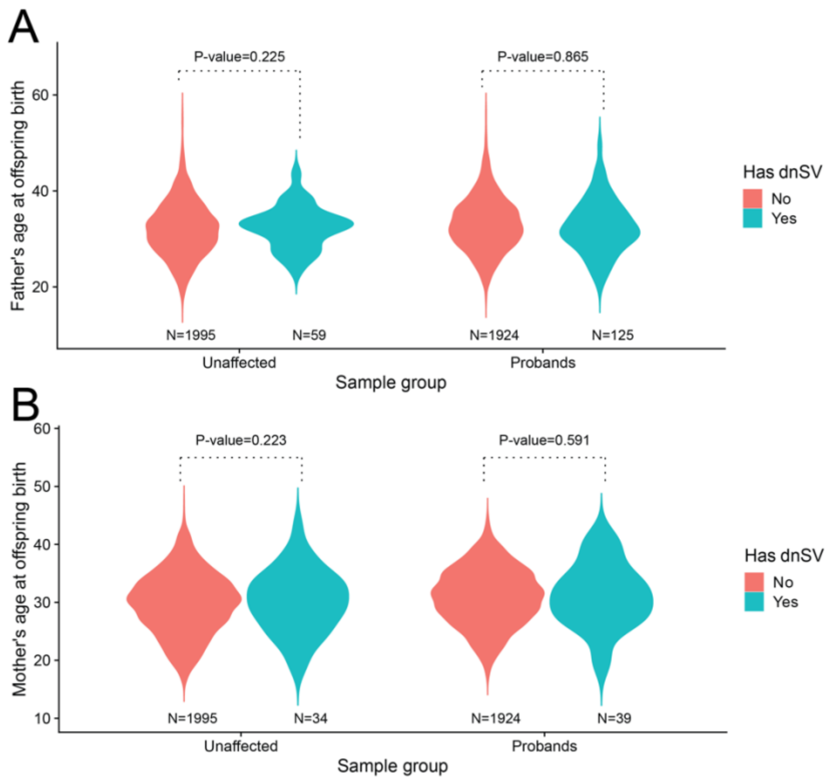


**Supplemental Figure 2. Correlations of parental age and de novo structural variant rate using phased variants. A.** One-sided Wilcoxon rank-sum test for an increase in father's age for samples with vs without at least one paternally derived dnSV. No significant difference in either siblings or probands. **B.** One-sided Wilcoxon rank-sum test for an increase in mother's age for samples with vs without at least one maternally derived dnSV. No significant difference in either siblings or probands.

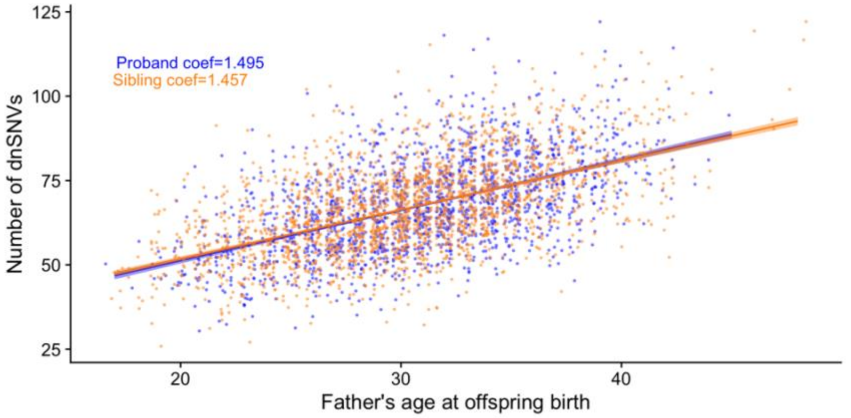


**Supplemental Figure 5. Correlation test between paternal age and *de novo* SNV count.** A Poisson regression was used to test the correlation of the count of de novo SNVs for most samples in the CEPH and SFARI cohorts with the paternal age.

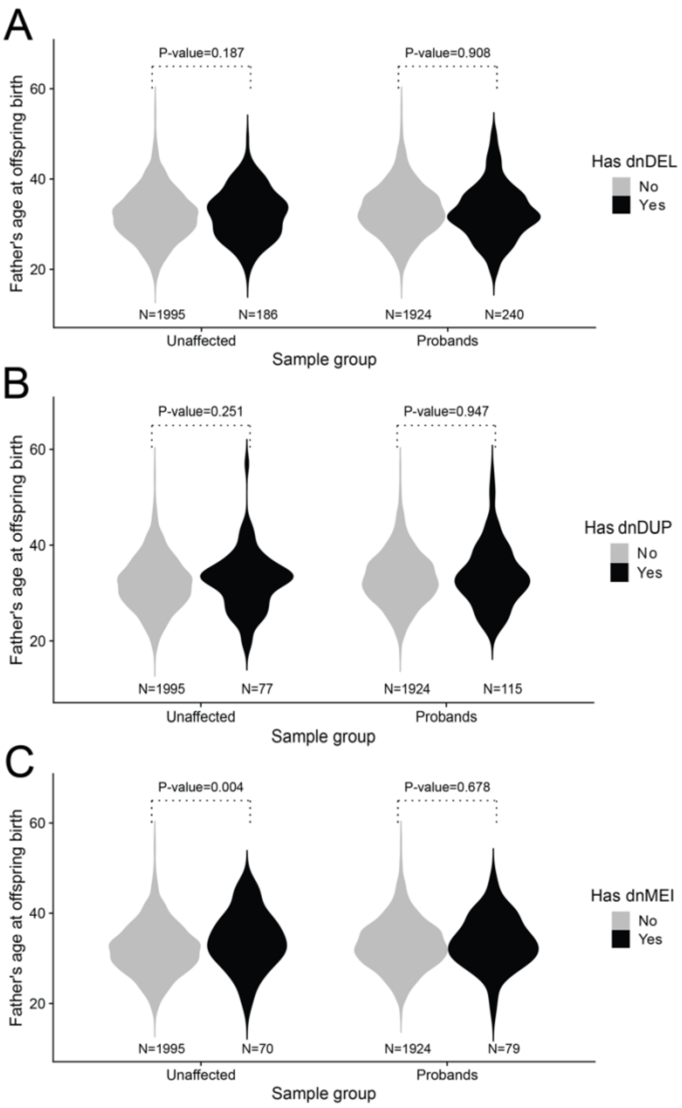


**Supplemental Figure 6. Correlations of paternal age and de novo structural variant rate by SV type. A.** One-sided Wilcoxon rank-sum test for an increase in father's age for samples with vs without at least one dnDEL. No significant difference in either siblings or probands. **B.** One-sided Wilcoxon rank-sum test for an increase in father's age for samples with vs without at least one dnDUP. No significant difference in either siblings or probands. **C.** One-sided Wilcoxon rank-sum test for an increase in father's age for samples with vs without at least one dnDEL. Fathers of offspring with dnMEIs are significantly older among siblings, but not among probands.
